# Opto4E-BP, an optogenetic tool for inducible, reversible, and cell type-specific inhibition of translation initiation

**DOI:** 10.1101/2023.08.30.554643

**Authors:** Jessica M. Alapin, Muhaned S. Mohamed, Prerana Shrestha, Houda G. Khaled, Anna G. Vorabyeva, Heather L. Bowling, Mauricio M. Oliveira, Eric Klann

## Abstract

The protein kinase mechanistic target of rapamycin complex 1 (mTORC1) is one of the primary triggers for initiating cap-dependent translation. Amongst its functions, mTORC1 phosphorylates eIF4E-binding proteins (4E-BPs), which prevents them from binding to eIF4E and thereby enables translation initiation. mTORC1 signaling is required for multiple forms of protein synthesis-dependent synaptic plasticity and various forms of long-term memory (LTM), including associative threat memory. However, the approaches used thus far to target mTORC1 and its effectors, such as pharmacological inhibitors or genetic knockouts, lack fine spatial and temporal control. The development of a conditional and inducible eIF4E knockdown mouse line partially solved the issue of spatial control, but still lacked optimal temporal control to study memory consolidation. Here, we have designed a novel optogenetic tool (Opto4E-BP) for cell type-specific, light-dependent regulation of eIF4E in the brain. We show that light-activation of Opto4E-BP decreases protein synthesis in HEK cells and primary mouse neurons. *In situ*, light-activation of Opto4E-BP in excitatory neurons decreased protein synthesis in acute amygdala slices. Finally, light activation of Opto4E-BP in principal excitatory neurons in the lateral amygdala (LA) of mice after training blocked the consolidation of LTM. The development of this novel optogenetic tool to modulate eIF4E-dependent translation with spatiotemporal precision will permit future studies to unravel the complex relationship between protein synthesis and the consolidation of LTM.

## Introduction

Beginning with the first experiments injecting the antibiotic and translation elongation inhibitor puromycin into mice, translation has been known to be required for the formation of long-term memory^1^. Since that groundbreaking study, many of the mechanisms of translation have been discovered, including initiation, ribosome recruitment, elongation, as well as the regulation of the translation factors involved in each of these steps. However, it was not until the mid-2000s that studies began to delineate the translational control pathways involved in both protein synthesis-dependent long-lasting synaptic plasticity and memory in the rodent brain. Much of this work has focused on translation initiation, which is the rate-limiting step in protein synthesis and where the majority of translational control takes place^2–5^.

The protein kinase mTOR, when in a complex with the adaptor protein Raptor (mTOR/Raptor is known as mTORC1), is a prime regulatory of cap-dependent translation initiation via phosphorylation of its substrates eIF4E-binding proteins (4E-BPs) and p70 S6 kinase 1 (S6K1)^5^. mTORC1 binds 4E-BPs (denoted as 4E-BP2, the brain-enriched isoform) and S6K1. mTORC1-dependent phosphorylation of 4E-BP2 and S6K1 is prevented by rapamycin, which disrupts the interaction of mTOR with Raptor. Unphosphorylated 4E-BP2 binds tightly to eIF4E to suppress translation initiation, whereas 4E-BP2 phosphorylated by mTORC1 does not bind eIF4E, which allows formation of the cap-dependent initiation complex eIF4F (eIF4E+ eIF4G+eIF4A1). mTORC1 also regulates translation initiation by phosphorylating S6K1, which then phosphorylates the downstream effectors eIF4B and PDCD4. The helicase activity of eIF4A1 is enhanced directly by phosphorylation of eIF4B and indirectly by the phosphorylation of PDCD4, which normally suppresses eIF4A1^6–8^. mTORC1 signaling is required for multiple forms of protein synthesis-dependent synaptic plasticity and various forms of long-term memory (LTM), including associative threat memory ^7–10^.

Previous studies have shown that the mTORC1 effectors eIF4E and S6K1 have specific roles in associative threat memory consolidation, reconsolidation, and extinction. For example, inhibition of eIF4F with the small molecule 4EGI-1, which inhibits eIF4E-eIF4G interactions^3^, blocked the consolidation, but not the reconsolidation, of associative threat memory in the rat amygdala^2^. In contrast, concomitant activation of eIF4E and S6K1 is required for the reconsolidation of associative threat memory in mice^11^. Notably, pharmacological inhibitors such as 4EGI-1 offer temporal control but lack cell type-specificity.

Recently, the generation of a mouse line for the conditional and inducible knockdown of eIF4E (4Ekd) mice was utilized to demonstrate that eIF4E-dependent translation in principal excitatory neurons of the lateral amygdala (LA) and somatostatin-positive inhibitory neurons of the centrolateral (CeL) amygdala is required for threat memory^12^ Moreover, this mouse line was used to show that eIF4E-dependent translation in protein kinase C δ-positive inhibitory neurons in the CeL is required for safety memory^13^. However, although the 4Ekd mice provide cell-type-specificity, they rely on a Tet-Off system to knockdown eIF4E, and therefore lack precise on-off control for investigating the temporal window for eIF4E-dependent translation in memory consolidation.

Viral-mediated gene transfer, combined with optogenetic approaches can allow for the cell type-specific expression of effector molecules in brain areas of interest, which can then be activated by light in matter of seconds, providing more precise spatiotemporal resolution^13–17^. Using this approach, we have now developed an optogenetic tool to achieve finer spatiotemporal resolution of eIF4E-dependent translation inhibition in neurons with a circularly permuted LOV inhibitor of protein synthesis (cLIPS ^17^. cLIPS is a first-generation light-switchable Opto4E-BP that rapidly binds to eIF4E and blocks cap-dependent translation by disrupting eIF4E-eIF4G interactions. In the present study, we show that light-activation of this Opto4E-BP effectively decreases protein synthesis in HEK293 cells and cultured neurons. Moreover, light-activation of Opto4E-BP in acute amygdala slices following in vivo viral expression in excitatory neurons resulted in decreased protein synthesis. We also show that light activation of Opto4E-BP in excitatory neurons of the LA 30 minutes after cued threat conditioning inhibits LTM consolidation in mice. Thus, this novel tool, Opto4E-BP, allows for detailed investigation of eIF4E-dependent protein synthesis with high spatiotemporal resolution in neuronal cells, in vitro, ex vivo, and in vivo.

## Results

### An optogenetic tool for cell-type specific protein synthesis inhibition

We have engineered a novel optogenetic tool to inhibit translation initiation in neurons, based on cLIPS^17^ a first-generation light-switchable Opto4E-BP that rapidly binds to eIF4E (Fig. 1a) to block cap-dependent translation by disrupting eIF4E-eIF4G interactions. This construct was first utilized in yeast cells^17^ and has now been adapted to allow expression in mammalian cells (HEK293 cells and neurons), using cell-type specific promoters. The 4E-BP2 fragment in cLIPS is comprised of the wild-type 4E-BP2 sequence from position 51-85 with mutated phosphorylation sites (S65A and T70A), whereas the photoswitchable domain in cLIPS is a circularly permuted version of *As*LOV2, the second light-oxygen-voltage domain of *Avena sativa* phototropin 1 ^17–22^, containing a point mutation V92I that slows the photocycle and thereby increases the light sensitivity^17^. To enable visualization of cells expressing Opto4E-BP, a fluorescence reporter mCherry was included in the construct and separated from Opto4E-BP with self-cleaving peptide P2A allowing ribosome reinitiation^23,24^ (Fig. 1b-c).

**Figure 1.**
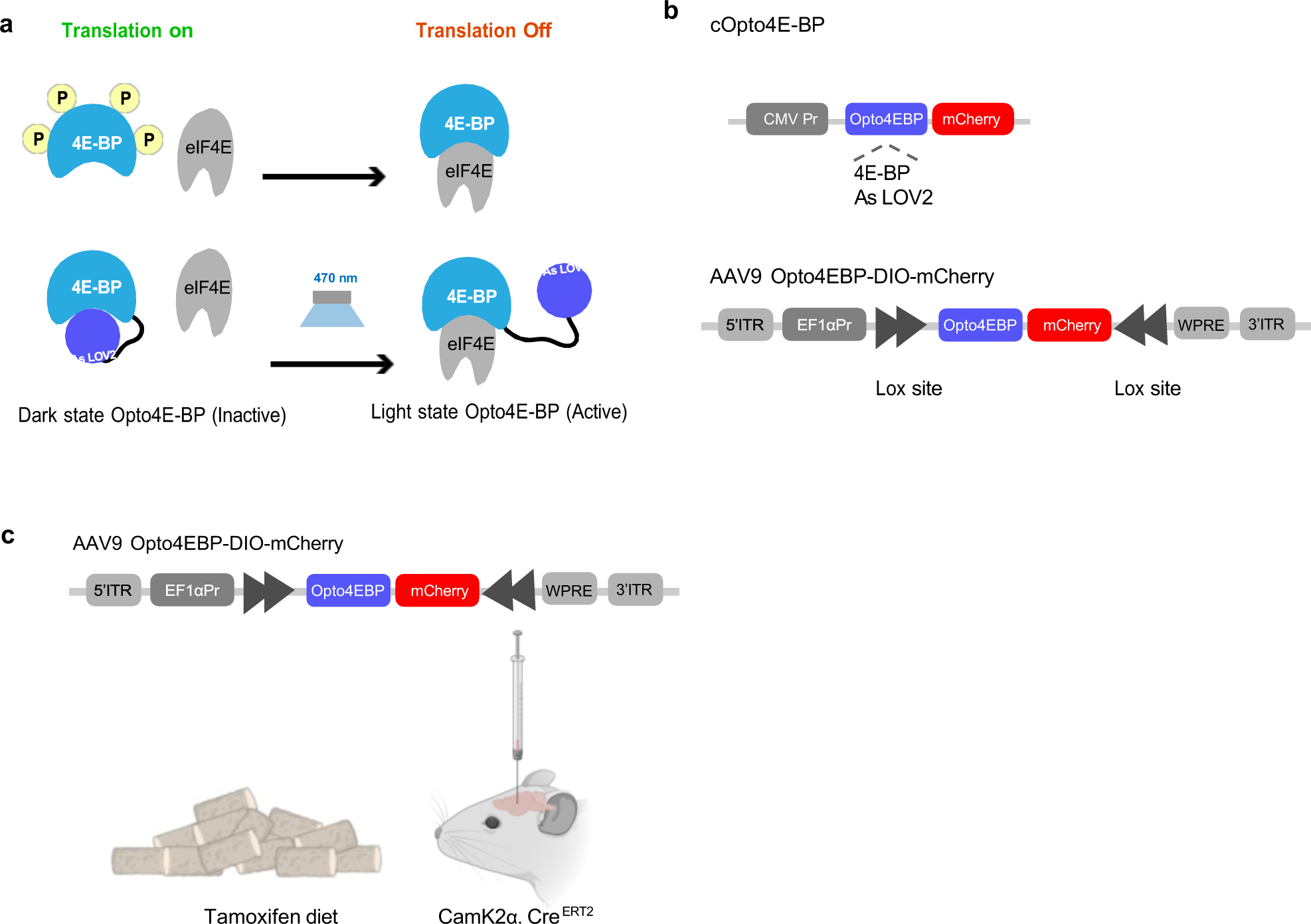
a. Diagram depicting the mechanism of Opto4E-BP activation. After blue light stimulation of Opto4E-BP and exposure of the binding site, 4E-BP binds to eIF4E to inhibit cap-dependent translation. b. Schematic representation of the cOpto4E-BP plasmid for cell culture transfection, and AAV9Opto4EBP-DIO-mCherry viral construct for neuronal expression. c. Tamoxifen inducible model for Opto4E-BP expression in vivo, in CamK2α.Cre^ERT2^ mice.

As a measure of translation, we performed <u>S</u>urface <u>S</u>ensing of <u>T</u>ranslation (SUnSET)^25^ in HEK293 cells with CMV.cLIPs using puromycin (Fig 2a). We found that blue light exposure (470 nm, ∼1.7 mW) for 15 minutes reduced basal and insulin-stimulated *de novo* protein synthesis (Fig. 2b). De novo protein synthesis in HEK cells exposed to blue light start to recover with 30 minutes with a full recovery within 6 hours to levels of *de novo* protein synthesis similar to cells that were not exposed to light (Fig. 2c, Supplemental Fig. 1c). Overall, these findings indicate that Opto4E-BP efficiently blocks translation in a transient manner.

**Figure 2.**
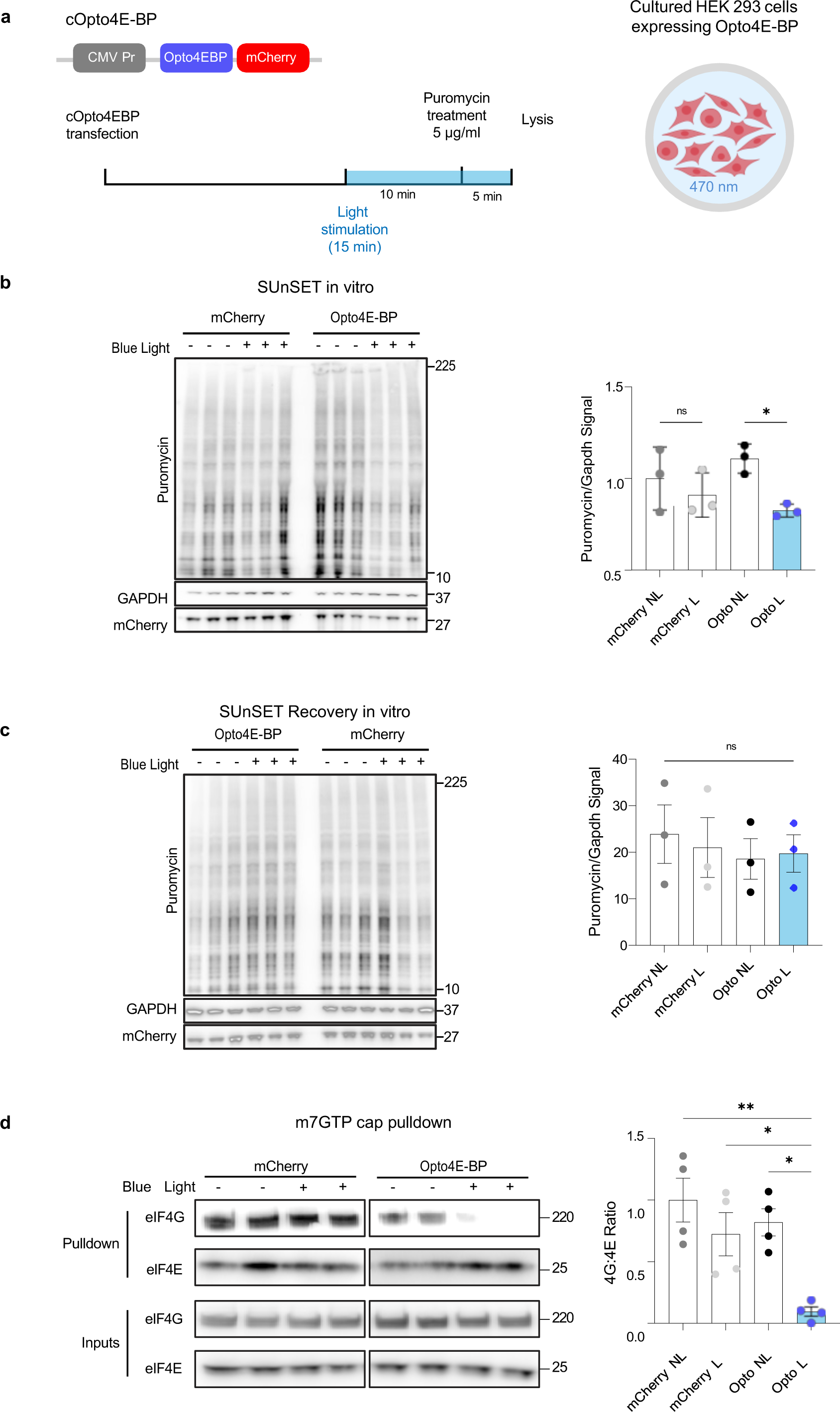
Blue light activation of cOpto4E-BP inhibits protein synthesis in HEK cells. a) HEK cells transfected with cOpto4E-BP or mCherry were given puromycin for 5 min in the presence (L) or absence (NL) f blue light. All samples were normalized to loading control GAPDH and no light. mCherry levels were measured to verify transfection efficiency. b) HEK cells exposed to blue light for 15 minutes, showed lower protein synthesis levels compared to the no light control (ANOVA multiple comparisons, Opto light versus no light, *p= 0.0318), One-way ANOVA F (3, 8) = 3412, p=0.0733. c) HEK cells exposed to blue light recovered within 6 hours to levels of *de novo* protein synthesis similar to cells that were not exposed to light, One-way ANOVA p=0.9064, F (3,8) =0.1809 d) Light activation of Opto4E-BP reduced eIF4E-eIF4G interactions, One-way ANOVA **p=0.0032 F (3,12) = 8.125 Data are presented as ± SEM.

In order to achieve optimal protein synthesis inhibition using Opto4E-BP, we transfected HEK293 cells with CMV.Opto4E-BP and performed different light treatment paradigms (Supplemental Fig. 1. a-b). Using SUnSET, we compared light treatment with blue light for 30 seconds versus 60 seconds and 2 cycles versus 10 cycles of light treatment (1 min on, 2 min off each cycle). In both experiments, we did not see a significant difference between the groups, and therefore we decided to use one consistent protocol (5 cycles) for subsequent experiments. In addition, Opto4E-BP is also capable of reducing insulin stimulated translation in HEK293 cells (Supplemental Fig. 1d). To confirm that activation of Opto4E-BP inhibited eIF4E-eIF4G interactions in HEK cells, we used a m^7^GTP cap pulldown assay following light/ no light stimulation of the cells. We found that light activation of Opto4E-BP reduced eIF4E-eIF4G interactions (Fig. 2d). These findings indicate that the light-activated Opto4E-BP prevents formation of eIF4F to prevent eIF4E-dependent translation.

### Generation of viral vectors expressing Opto4E-BP to inhibit protein synthesis in cultured neurons and brain slices

We next proceeded to investigate the efficacy of Opto4E-BP in neurons, with the ultimate goal of using it in vivo for studies of memory. We designed an adeno-associated viral (AAV) construct to express Opto4E-BP in neurons in a Cre-dependent manner (using a double inverted open reading frame) and included an mCherry tag to visualize expression (AAV9-EF1a-DIO-Opto4E-BP-mCherry) (Fig. 3a). We first tested the effectiveness of the virally-expressed Opto4E-BP by transducing primary neurons derived from BAFF53-Cre mice^26^ for pan-neuronal Cre expression. As a control, we used a virus driving Cre-dependent expression of mCherry only. At DIV 10, we performed light stimulation and labeled the *de novo* proteome using the puromycilation SUnSET assay, as described above. We observed a significant decrease in puromycilated proteins after exposure to light when neurons were transduced with AAV-Opto4E-BP, an effect that was not observed in neurons transduced with control virus and in no light-treated samples (Fig. 3b). Because western blots do not allow cell type-specific visualization of puromycilation, we also performed SUnSET using immunofluorescence, and confirmed this decrease is specific to Opto4E-BP-harboring neurons (Fig. 3c).

**Figure 3.**
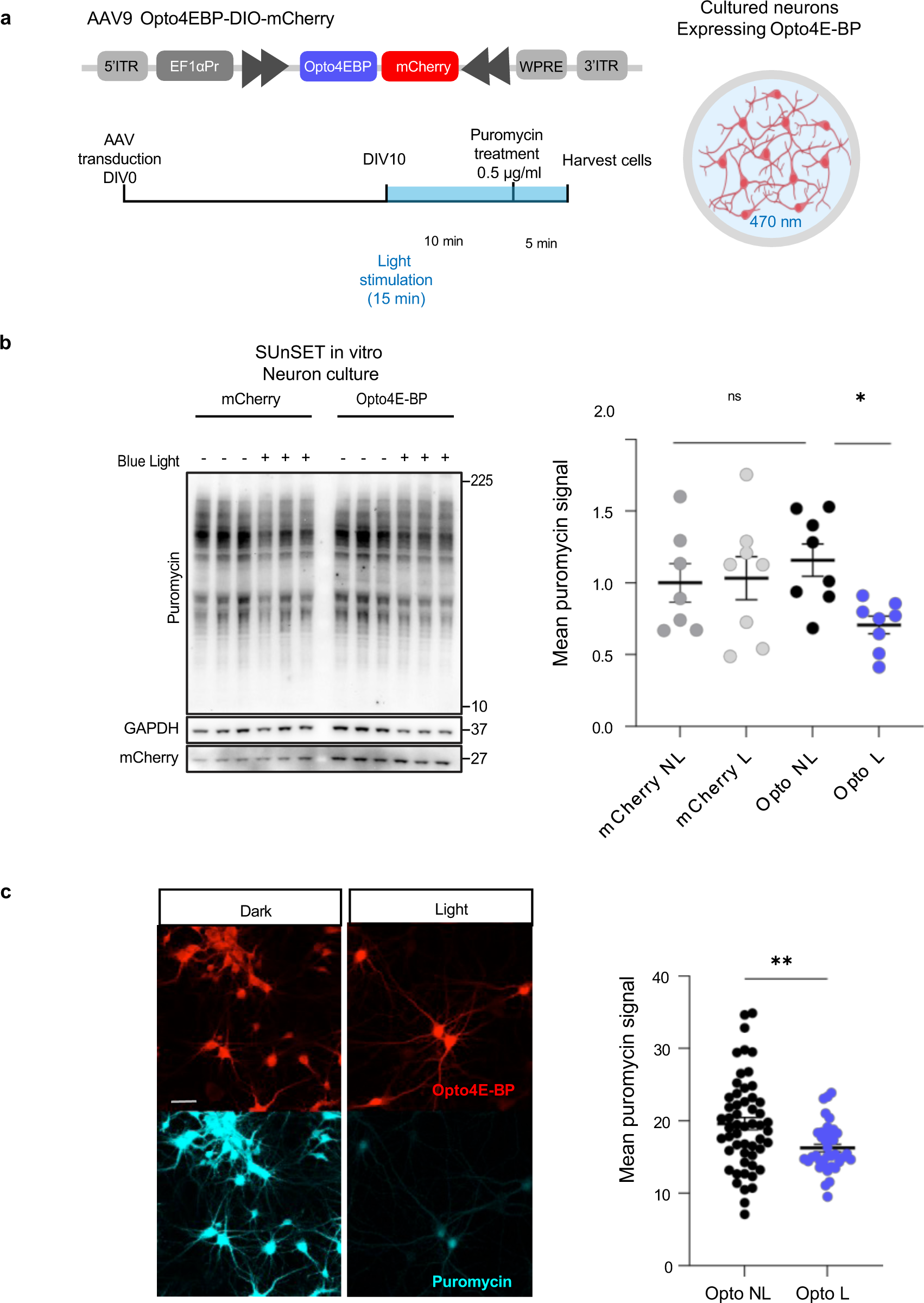
Blue light activation of AAV Opto4E-BP inhibits protein synthesis in cultured neurons. a) Light stimulation protocol and puromycin treatment. b) Primary neurons with Opto4E-BP/ mCherry controls were treated with puromycin labeling for 5 minutes in the presence (L) or absence (NL) of blue light. All samples were normalized to loading control GAPDH and no light. One-way ANOVA p=0.0665, ANOVA multiple comparisons L vs NL *p=0.0497 t-test (L vs NL**p=0.0032). c) Imaging of primary neurons transduced with Opto4E-BP treated with puromycin labeling for 5 minutes in the presence (L) or absence of blue light. Unpaired Student t-test**p=0.0030. Scale bar, 50 µm. Data are presented as ± SEM.

Given our interest in studying the role of *de novo* protein synthesis in the consolidation of emotional memories, we proceeded to conduct experiments in the lateral amygdala (LA), a brain structure, required for the consolidation of threat memories^27–29^. We first tested the efficiency of Opto4E-BP in blocking protein synthesis ex vivo using amygdala slices from CamK2α.Cre ^ERT2^ mice^30^ in order to express Opto4E-BP in excitatory neurons. Slices underwent metabolic labeling with l-azidohomoalinine (AHA) for visualization of newly synthesized proteins with fluorescent noncanonical amino acid tagging (FUNCAT)^31^ 45 minutes following AHA labeling, slices received 5 cycles of light treatment or no light (470 nm,1.7 mW, 1 min ON, 2 min OFF) (Fig 4a). We found that light-stimulated amygdala slices exhibited a significant reduction in *de novo* translation compared to no light controls (Fig. 4b). Thus, light stimulation of excitatory neurons in the LA of mice expressing Opto4E-BP, effectively inhibits *de novo* protein synthesis. Moreover, these findings demonstrate that Opto4E-BP can be expressed in vivo and can efficiently be used to control protein synthesis in ex vivo slice preparations.

**Figure 4.**
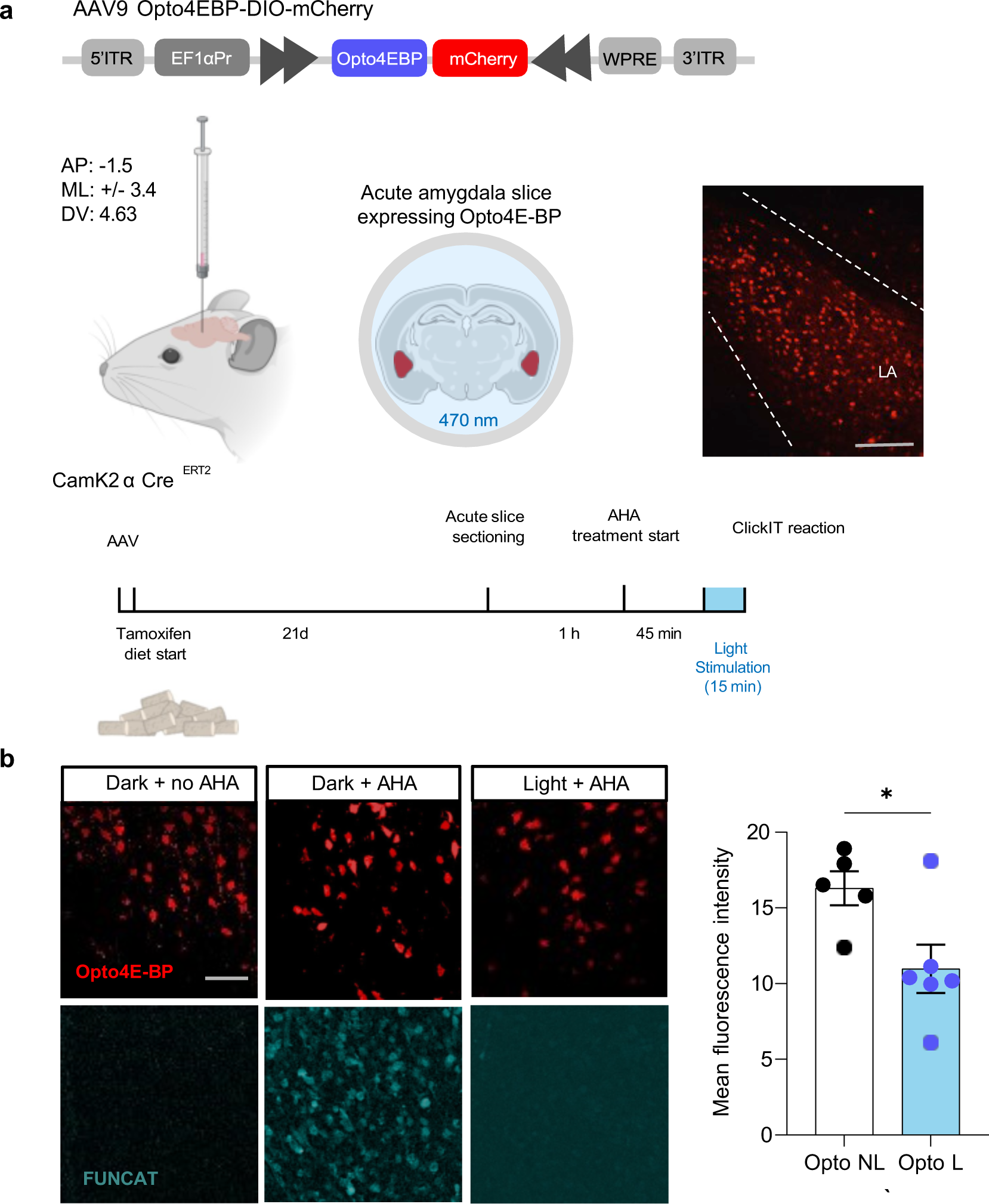
Blue light activation of AAV Opto4E-BP inhibits protein synthesis in acute amygdala slices following in vivo expression. a) Viral expression in the amygdala and light stimulation protocol. Scale bar, 200 µm. b-c) Acute amygdala slices expressing Opto4E-BP were labeled with AHA for 1 hour and exposed to blue light in the last 15 mins. Unpaired t-test *p=0.0280. Scale bar 50, µm. Data are presented as± SEM.

### Light activation of Opto4E-BP in the LA after cued threat conditioning inhibits LTM consolidation in mice

We proceeded to test the AAV9-EF1a-DIO-Opto4EBP-mCherry viral construct in vivo. We utilized this Opto4E-BP tool to determine temporal windows in excitatory neurons in the LA that require eIF4E-dependent protein synthesis for associative threat memory consolidation^32–34^. CamK2α. Cre ^ERT2^ mice were injected in the LA with either AAVOpto4E-BP or mCherry control virus. Expression of both viral constructs were found to be equal in Camkii positive neurons in the lateral amygdala (Supplemental Fig. 2). Immediately after the injection, mice were placed on a tamoxifen diet to induce expression of Cre. Four weeks after the injection, mice underwent cued threat conditioning training, and 30 minutes after training, they received either light stimulation, or no light, through optical cannulas in the amygdala (Fig 5.a). Light stimulation consisted of 5 cycles (470nm, 1.7 mW/mm^2^, 1 min ON, 2 min OFF) for 15 minutes (Fig 5b). We found that mice that received light treatment 30 minutes after training exhibited significantly lower levels of freezing during testing 24 hours later compared to all the control groups (4E-BP no light, mCherry light/ no light) (Fig 5c-e). These results are consistent with previous findings in the field regarding the temporal windows of *de novo* protein synthesis required for memory consolidation and show for the first time that eIF4E-mediated protein synthesis in excitatory neurons in the LA is required for threat memory consolidation^13,32–34^

**Figure 5.**
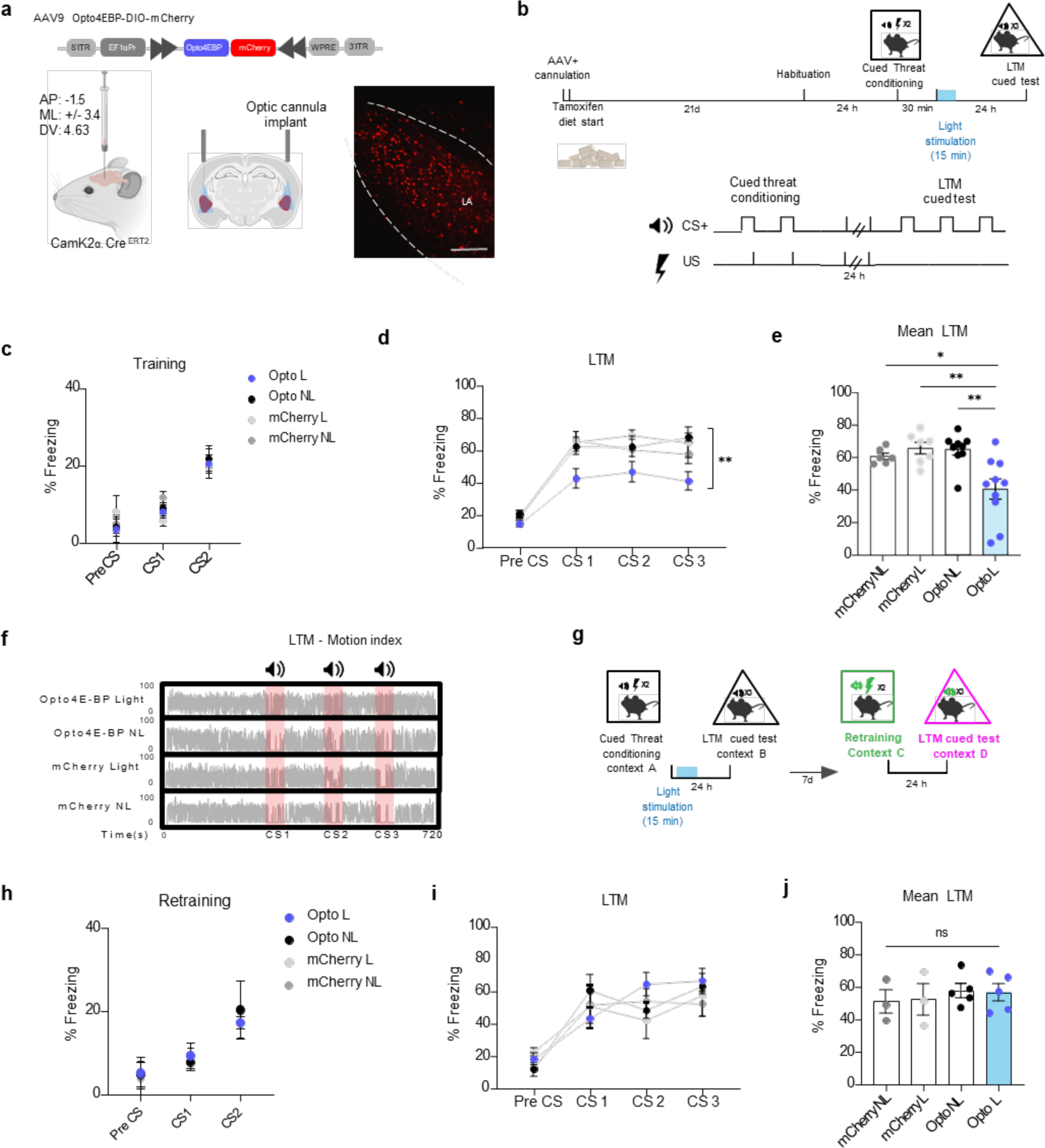
Blue light activation of AAV Opto4E-BP 30 minutes after cued threat conditioning inhibits LTM formation in mice. a) Viral expression in the amygdala and light stimulation protocol. CamK2α.Cre^ERT2^ mice infected with AAV Opto4E-BP/ mCherry controls in the amygdala underwent cued threat fear conditioning. b) Behavioral paradigm for cued threat conditioning with light stimulation. 30 minutes after training they received light stimulation/ no light in the amygdala for 15 minutes. 24 hours after training, mice were tested for cued threat LTM. c) Mice learned the association between CS and US during training. RM Two-way ANOVA with Bonferroni’s post-hoc test. CS: F (3,40) = 1.014, p=0.3964. d) Cued LTM was impaired Opto4E-BP light treated mice compared to controls across all three CS presentations. n=6-10 per group; RM Two-way ANOVA with Bonferroni’s post-hoc test. F (3, 26) = 5.289, **p=0.0024. e) Mean LTM was significantly impaired in Opto4E-BP light treated mice compared to controls F (3, 28) = 7.3411 (***p=0.0003) n = 6-10 per group; Unpaired t-test. Data are presented as ±SEM. f) Representative motion index of mice during Cued LTM test. g) Behavioral paradigm for cued threat conditioning for retraining mice a week after light stimulation. h) Mice learned the association between CS and US during training. RM Two-way ANOVA with Bonferroni’s post-hoc test. CS: F (3, 16) = 0.02418 p=0.9947. i) Cued LTM was intact in Opto4E-BP light treated mice compared to controls across all three CS presentations. n=3-5 per group; RM Two-way ANOVA with Bonferroni’s post-hoc test. F (3, 48) = 0.1771, p=0.9114 ns. j) Mean Cued LTM in Opto4E-BP light treated mice was not significantly different compared to controls. F (3, 12) = 0.2641 (ns p=0.8500) n = 3-5 per group; One-way ANOVA. Scale bar 200 µm. Data are presented as ± SEM.

In addition to testing light stimulation of Opto4E-BP 30 minutes after training, we performed experiments with light stimulation immediately after training, and found a reduction in the freezing response during testing 24 hours later (Supplemental Fig. 3). We also activated Opto4E-BP 5 hours after training, which is likely outside of the protein synthesis-dependent window required for memory consolidation ^32–34^. During the LTM test, mice in the Opto4E-BP light-treated group, exhibited normal levels of freezing similar to freezing in the no light control group (Supplemental Fig 4). We next tested whether Opto4E-BP activation would affect short-term memory formation, which is known to be protein synthesis-independent. Mice received light/no light treatment 30 minutes after threat conditioning and were tested 2 hours after training. We did not observe any difference in the freezing response between the groups (Supplemental Fig. 5). Because we found that the impact of Opto4E-BP activation on protein synthesis is rapidly reversed in vitro (Supplemental Fig. 2c), we determined whether the memory of mice that underwent threat conditioning would have intact memory at a later time point. Mice were retrained in a different context one week after receiving light stimulation. Notably, light-treated Opto4E-BP expressing mice that showed memory impairment in the first training session had intact memory, indicating that the transient blockade of protein synthesis is not sufficient to create lasting defects that impair consolidation (Fig 5g-j). Taken together, these findings indicate that Opto4E-BP can efficiently be used to prevent LTM consolidation in mice that received light treatment in the amygdala after training, specifically during the time window required for memory consolidation.

## Discussion

Translational control at the initiation step is paramount for proper cell function. There are a few chemogenetic tools available that allow one to study translation initiation in a cell type-specific and inducible manner, but the tools to study eIF4E-dependent translation lack fine temporal resolution^12,13^. To solve this problem, we developed and characterized a novel optogenetic tool, Opto4E-BP, for inducible and light-dependent regulation of eIF4E to probe the role of eIF4E-dependent protein synthesis in memory consolidation in vivo with extremely high spatiotemporal resolution. The present study shows that light-activation of Opto4E-BP effectively decreases protein synthesis in HEK293 cells and cultured neurons. Moreover, light-activation of Opto4E-BP in acute amygdala slices following in vivo viral expression in excitatory neurons resulted in decreased protein synthesis. We also show that light activation of Opto4E-BP after auditory threat conditioning inhibits LTM consolidation in mice that received light treatment 30 minutes after training, a crucial temporal window for LTM consolidation that requires *de novo* protein synthesis. Moreover, when mice expressing Opto4E-BP were retrained in a different context one week after light stimulation, they exhibited normal LTM formation, indicating that activation of Opto4E-BP can be reversed in vivo and does not damage the neuronal circuitry in which it is expressed. These results are consistent with a critical role for eIF4E-dependent protein synthesis in threat memory consolidation^13^.

We anticipate that the application of Opto4E-BP will be useful to address a number of outstanding questions concerning the role of local protein synthesis in both long-lasting synaptic plasticity and memory consolidation. For example, Opto4E-BP could be helpful for studying the requirement of local axonal translation in memory consolidation^13,35–40^ as it can be expressed in excitatory neurons in either the thalamus or the auditory cortex that send projections to the LA in order to determine whether eIF4E-dependent translation LA axons is required for memory consolidation.

In our study, we demonstrated that Opto4E-BP can be used for investigating protein synthesis in cell culture and can help determine the temporal windows and cell types that require eIF4E-dependent protein synthesis for associative memory consolidation in vivo. In addition to excitatory neurons, Opto4E-BP has great potential for a variety of studies to study eIF4E-dependent translation in other brain cells, including inhibitory neurons, astrocytes, and microglia, as well as cells in peripheral tissues. The versatility of Opto4E-BP allows for its application in studies far beyond the study of memory formation. For example, dysfunctional mTORC1 signaling has been shown to play a role in several types of autism spectrum disorder (ASD), and was found to alter neuron structure and function, leading to focal malformation of cortical development (FMCD) and intractable epilepsy^41–45^. eIF4E has been found to be important for maintenance of circadian rhythms^46–50^ and outside of the brain, has been shown to play a role in cancer development ^51–54^.

In conclusion, we have developed a novel genetically encoded cell-type specific optogenetic tool with high spatiotemporal resolution to inhibit eIF4E-dependent translation in neuronal cells in vitro, ex vivo, and in vivo. Opto4E-BP is a versatile and efficient tool to inhibit cap-dependent protein synthesis, which can be used in studies of various memory processes, models of ASD and FMCD, and in other research areas outside of the field of neuroscience.

## Materials and Methods

### Drugs and chemicals

Puromycin (Sigma, P8833) was dissolved in ddH2O at 25 µg/µl, and this stock was freshly diluted for SUnSET assays in vitro. FUNCAT ex vivo *experiments* were performed using 100 mM azidohomoalanine (AHA) (Fisher, NC0667352) dissolved in distilled water. AHA was added to acute amygdala slices submerged in ACSF (25 mM glucose, 125 mM NaCl, 2.5 mM KCl, 1 mM MgCl2, 2 mM CaCl2, 1.25 mM NaH2PO4, and 25 mM NaHCO3). Tamoxifen was administered via the rodent chow at 40 mg/kg (Bio-Serv, F4159). CamK2α.Cre ^ERT2^ mice were fed this tamoxifen diet, ad libitum, immediately after surgery for four weeks until the end of the behavioral tasks.

### Cloning of cOpto4E-BP plasmid

The cLIPs construct was cloned into a pcDNA3.1 backbone with a CMV promoter (Addgene). Cloning was performed as previously described^13^. All constructs were verified using sequencing.

### HEK293 cell culture and transfection

HEK293 cells were grown in DMEM media (10% FBS, 1x P/S, 1x L-glutamine) at 37 °C, 5% CO2, and split into 12-well plates at a density of 250K cells/well. Cells were transfected the next day with plasmids encoding CMV-Opto-4EBP-mCherry or CMV-mCherry using lipofectamine 3000 according to the protocol of the manufacturer, and kept in the dark at all times prior to blue light exposure to avoid Opto4E-BP activation. Media was changed 18-24 hours after transfection, 1-2 hours before blue light exposure. A blue light LED array (Onstate Technologies) was placed inside the cell incubator above the 12-well plate and connected to a plug-in mechanical timer to customize the light on/off cycle. Wells not receiving light exposure were covered with aluminum foil. Puromycin (5 µg/mL) was added to wells during a dark cycle in the last 5 minutes of light cycle exposure. Afterward, cells were immediately lysed by removing media and adding RIPA buffer to wells (1x Protease Inhibitory Cocktail, 1x EDTA). BCA protein assay (Pierce) was used to measure protein concentrations.

### In vitro m^7^GTP cap pulldown experiments

The m^7^GTP cap pulldown experiments were performed as previously described^53^. HEK293 Cells were grown to 60% confluency in 10cm dishes and transfected with 20 µg of mCherry or Opto4E-BP construct using lipofectamine 3000 (Thermo Fisher). Cells then were treated with 470 nm blue light for 5 cycles (1 min, 2 min off) for 15 minutes. Cells then were scraped in ice cold PBS, spun at 300g for 5 minutes and the pellets immediately lysed in pull down lysis buffer (150 mM NaCl, 10 mM MgCl_2_, 30 mM Tris buffer (pH 8.0), 1 mM DTT, 1.5% Triton X-100, 1x Halt Protease and phosphatase inhibitor cocktail (Thermo Fisher). Protein concentrations were measured using BCA and 300 µg of protein lysate was added to 50 µl of m^7^GTP beads (Jena Bioscience) for 2h at 4°C. The light-treated group was continually treated with 470 nm light for the duration of the incubation. Beads then were centrifuged at 6000 rpm for 1 minute and then washed 3 times with wash buffer (100 mM KCl, 50 mM Tris buffer (pH 7.4), 5 mM MgCl_2_, 0.5% Triton X-100). The beads then were eluted in 50 µl 1x Laemmli buffer supplemented with 25 mM DTT.

### Viral vectors for neuronal expression

AAV9-hsyn-DIO-mCherry (1x 10^13^ GC/mL titer) was purchased from Adgene (#50459). AAV9-EF1a-DIO-Opto4EBP-mCherry (2.1x 10^13^ GC/mL titer) was custom ordered from Vector Biolabs.

### Primary neuron cultures and transduction

Primary neurons derived from E16 pups were plated at 2.5×10^5^ cells per well for imaging or 2×10^6^ cells per well for western blotting. Neurons were transduced at DIV 1 with mCherry or Opto4E-BP viral vector using 1000 genomic copies per cell. At DIV 10, cells were stimulated with 488nm for 5 cycles (1 min, 2 min off) for 15 minutes. After the 3rd cycle (10 minutes into stimulation), cells were treated with 5 µg/ml of puromycin. After 5 minutes, cells were immediately lysed in RIPA Buffer, and lysates were used for either western blots or fixed with 4% PFA for immunocytochemistry.

### Immunocytochemistry

Immunocytochemistry in primary neurons was performed by permeabilizing with 0.2% Triton X followed by blocking for 1 hour with 5% normal goat serum. Neurons were then incubated with the following primary antibodies in blocking buffer overnight guinea pig MAP2 (1:1000) (Synaptic Systems), rat mCherry (1:1000) (Invitrogen), mouse puromycin (1:1000) (Millipore). Neurons were washed 3 times with PBS with 0.1% Tween-20 and were incubated for 1 hour in the following secondary antibodies diluted in blocking buffer anti-guinea pig AF 405 (1:750), anti-rat AF 594 (1:750), and anti-mouse AF 647 (1:750) (Invitrogen). Coverslips were mounted on slides using Prolong-Gold (Invitrogen) and imaged using a confocal (Leica SP5) using a 40x oil immersion objective.

### Animals

All experiments were performed according to the guidelines of the National Institutes of Health (NIH) and were approved by the University Animal Welfare Committee of New York University. Neuronal culture experiments were performed using BAFF53-Cre mice^26^ (with pan-neuronal Cre expression). For all acute slice and behavioral experiments, 12-week-old CamK2α.Cre ^ERT2^ mice were used, in which a tamoxifen-inducible Cre is expressed in CamK2α neurons^54^. Both female and male mice were used for all experiments; no sex-specific differences were noted. Mice were kept at stable conditions (78°F temperature, on a 12h/12h dark/light cycle, humidity of ∼ 45%) and received food and water ad libitum.

### Stereotaxic surgeries

For experiments performed in ex vivo slices and behavior, CamK2α.Cre^ERT2^ mice were anesthetized via isoflurane inhalation using an automated system (SomnoSuite, from Kent Scientific). Mice then were bilaterally injected with either mCherry control or DIO-Opto4EBP-mCherry in the lateral amygdala (using a Neurostar robotic injection system). Coordinates for LA (−1.5 AP, +/-3.5 ML, 4.6 DV). Injection rate: 0.04 µl/min 200 nanoliters per hemisphere. For behavioral experiments, optic cannulas were implanted above the LA immediately after the AAV injection. Coordinates for cannulation (AP: −1.5, ML:3.5, DV:4.1) (Thor labs).

### Metabolic labeling with AHA and ex vivo FUNCAT

CamK2α.Cre ^ERT2^ mice expressing Opto4E-BP, were sacrificed by cervical dislocation, and 300 µm ex vivo brain slices containing the amygdala were prepared using the aforementioned conditions. Slices were maintained in 1:1 carboxygenated ACSF: Cutting solution (25 mM glucose, 125 mM NaCl, 2.5 mM KCl, 1 mM MgCl_2_,2 mM,2 mM CaCl_2_, 1.25 mM NaH_2_PO_4_, and 25 mM NaHCO_3_), and allowed to recover for 1 hour. Slices then were treated with 100 mM AHA For 1 hour. 45 minutes into the AHA treatment, slices received light stimulation for 15 minutes (470 nm,1.7 mW/mm^2^, 1 min, 2 min off). Slices then were fixed with 4% PFA. 24 hours after fixation, slices underwent protein synthesis labeling using the Click-iT Cell Reaction Buffer Kit (Invitrogen).

### Cued threat conditioning and optogenetics in vivo

Behavioral experiments were performed during light cycles. Mice were divided randomly into the different experimental conditions and were tested in a random order for any given experimental paradigm. Behavioral data was collected using Freeze Frame 4 software (ActiMetrics), and the experimenters were blind to treatment conditions. Mice were first habituated for 15 min to the fear conditioning chamber (Colbourn instruments) and then habituated 15 min to the optogenetic setup (Thor labs). On the second day, mice underwent cued threat conditioning (cTC) training that consisted of 2 CS-US presentations (0.5mA foot shock, 85dB, 5kHz pure tone). The inter-trial interval (ITI) was 2 minutes and after the second tone-shock presentation, mice remained in the chamber for an additional 120 seconds. Immediately after training, mice received light stimulation with 5 cycles (470 nm,1.7 mW/mm^2^, 1 min ON, 2 min OFF) for 15 minutes via the optic cannulas (Thor labs). 24 hours after training, mice underwent cued threat memory testing in a different context (red light with a plexiglass platform and vanilla-scented bedding) and were presented 3 times with the same tone. Additional experiments were performed where different groups of mice received light stimulation for either 30 minutes or 5 hours after training. Only mice that had viral expression in the amygdala on both hemispheres were included in the data analysis. In the case of mice that received light stimulation 30 minutes after training, they were retrained and tested 7 days later under different conditions (1 kHz tone and peppermint-scented rubber mat during the LTM test).

### Image analysis

Imaging data from immunocytochemistry experiments was acquired using an SP5 confocal microscope (Leica) with 20× objective lens (with 1× zoom) and z-stacks (approximately ten optical sections with 0.563-μm step size) for three coronal sections per mouse from AP −1.4 mm to −1.70 mm (n = 3 mice) were collected. The imaging data was analyzed with ImageJ using the Bio-Formats importer plugin. A maximum projection of the z-stacks was generated, followed by manual outlining of individual cells and mean fluorescence intensity measurements using the drawing and measure tools. Mean fluorescence intensity values for all cell measurements were normalized to the mean fluorescence intensity for controls.

### Statistical analysis

Sample sizes were determined similarly to reports in previous publications^12,13^. Statistical analysis was performed using GraphPad Prism 8 (GraphPad Software). Data are shown as mean ± SEM. Normal distribution of data was assumed. Multiple group comparisons were analyzed using One-Way ANOVA or Repeated Measures Two-way ANOVA. Data from the comparison between two groups was performed using unpaired Student’s t-test. α level of 0.05. p-value was used for analysis.

## Acknowledgements

This work was supported by NIH grants NS047384 and NS122316 (E.K.). We would like to thank Professor Andrew Woolley for kindly providing the Opto4E-BP plasmid construct. We would also like to thank Lilly Cole and Sarah Reyman for assistance with HEK293 experiments, and Natalie Ito for assisting with stereotactic surgeries and histology.

## Author contributions

E.K. J.M.A and M.S.M, co-wrote the manuscript. J.M.A led the project under the supervision of E.K. E.K., J.M.A, M.S.M and P.S. designed the experiments. M.S.M, H.G.K and H.B, performed the in vitro experiments in HEK cells and cultured neurons. J.M.A. Performed the behavioral experiments and ex vivo slice experiments. All authors contributed to the discussions and preparation of manuscript.

## Notes

### Competing Interest Statement

The authors have declared no competing interest.

