## Supplemental Figures for "Opto4E-BP, an optogenetic tool for inducible, reversible, and cell type-specific inhibition of translation initiation"

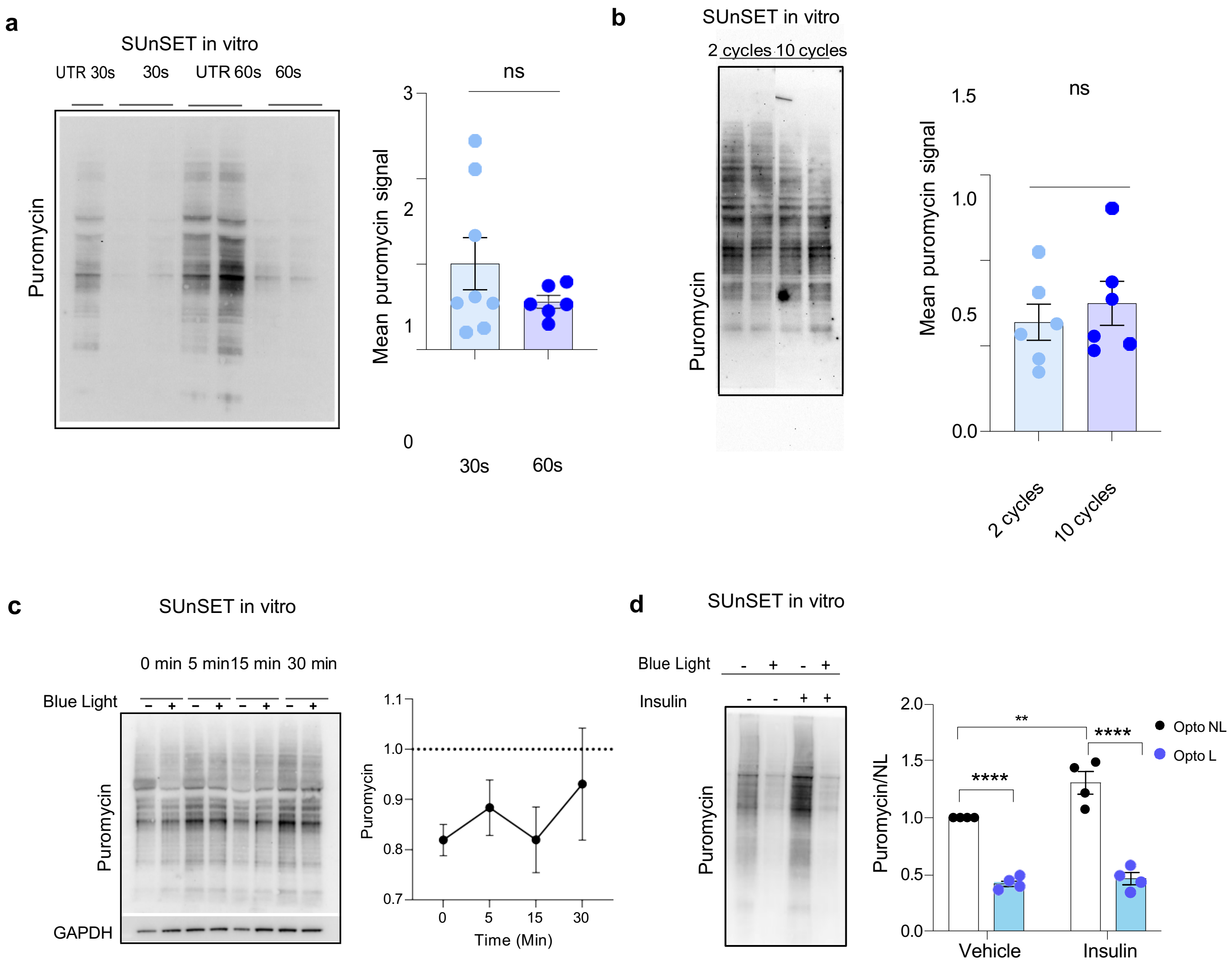


**Supplemental Fig. 1.** Blue light treatment optimization of cOpto4E-BP in HEK cell culture. a) Puromycilation assay of cells expressing cOpto4E-BP treated with blue light 30 secs vs 60 secs showed no difference between the two groups. Unpaired t-test ns (p=0.2376). b) Puromycilation assay of cells expressing cOpto4E-BP treated with blue light for 2 cycles vs 10 cycles showed no difference between the two groups. Unpaired t-test ns (p=0.5198). c) HEK cells transfected with cOpto4E-BP, treated with blue light, recover to normal levels of protein synthesis within 30 minutes. d) HEK cells transfected with cOpto4E-BP were given puromycin for 2 hrs in the presence (L) or absence (NL) of blue light with and without insulin (85 µM). mCherry levels were measured to verify transfection efficiency. Samples were normalized to total protein and no light, no insulin. Two-way ANOVA ****p<0.0001, **p<0.01, ns = not significant. Data are presented as ± SEM.

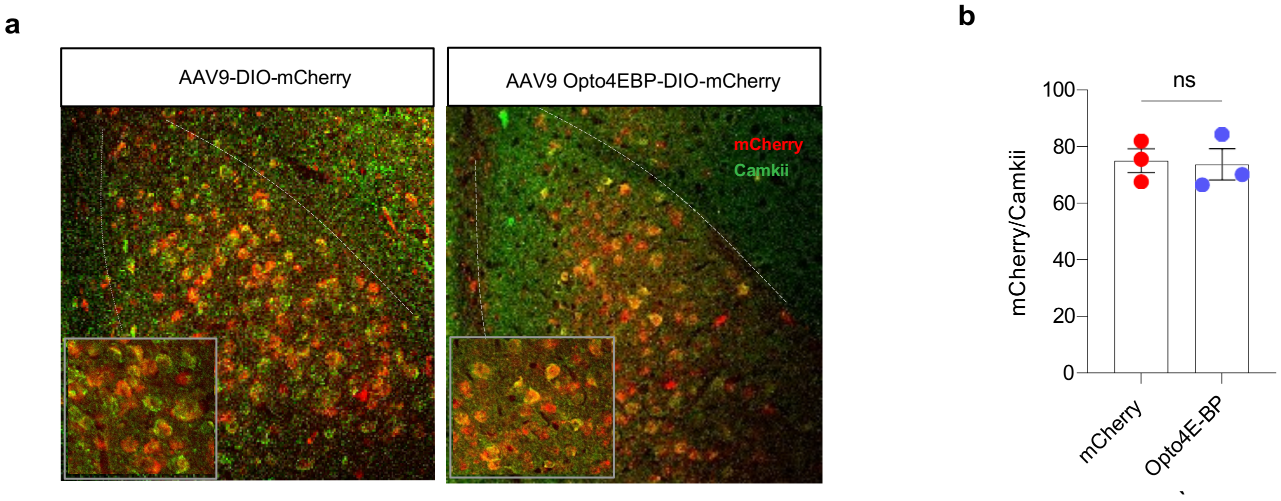


**Supplemental Fig. 2.** Viral expression quantification in the LA of CamK2α.Cre $^{ERT2}$ mice injected with AAV Opto4E-BP or AAV mCherry. No difference between the two groups. Unpaired Student's t-test p=0.8556 ns=not significant. Data are presented as ± SEM.

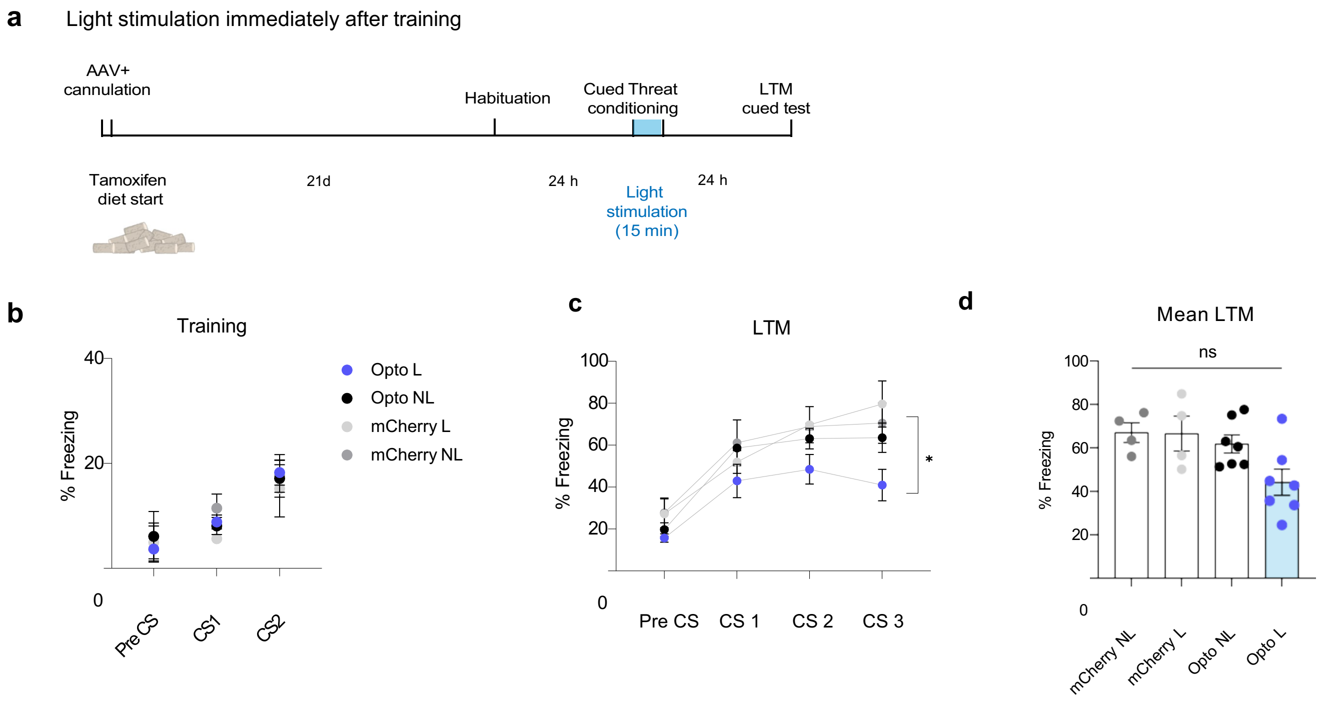


**Supplemental Fig. 3**. Blue light activation of AAV Opto4E-BP immediately after cued threat conditioning inhibits LTM formation in mice. a) Light stimulation protocol. CamK2α.Cre $^{ERT2}$ mice infected with AAV Opto4E-BP or mCherry controls in the amygdala underwent cued threat fear conditioning. Immediately after training they received light stimulation/ no light in the amygdala for 15 minutes. 24 hours after training, mice were tested for cued threat LTM. b) Behavioral paradigm for cued threat conditioning with light stimulation c) Mice learned the association between CS and US during training. RM Two-way ANOVA with Bonferroni's post-hoc test. CS: $F(3,54) = 0.4618$, $p=0.712$. c) Cued LTM was partially impaired Opto4E-BP light treated mice compared to controls across all three CS presentations. n=4-7 per group; RM Two-way ANOVA with Bonferroni's post-hoc test. $F(3,54) = 7.232$, $^{*}p=0.0271$. d) Mean cTC LTM was significantly impaired in Opto4E-BP light treated mice compared to controls One-way ANOVA $F(3, 18) = 3.788$ ($^{*}p=0.0288$) n = 4-7 per group, multiple comparisons showed no significant differences between the groups. Data are presented as ± SEM.

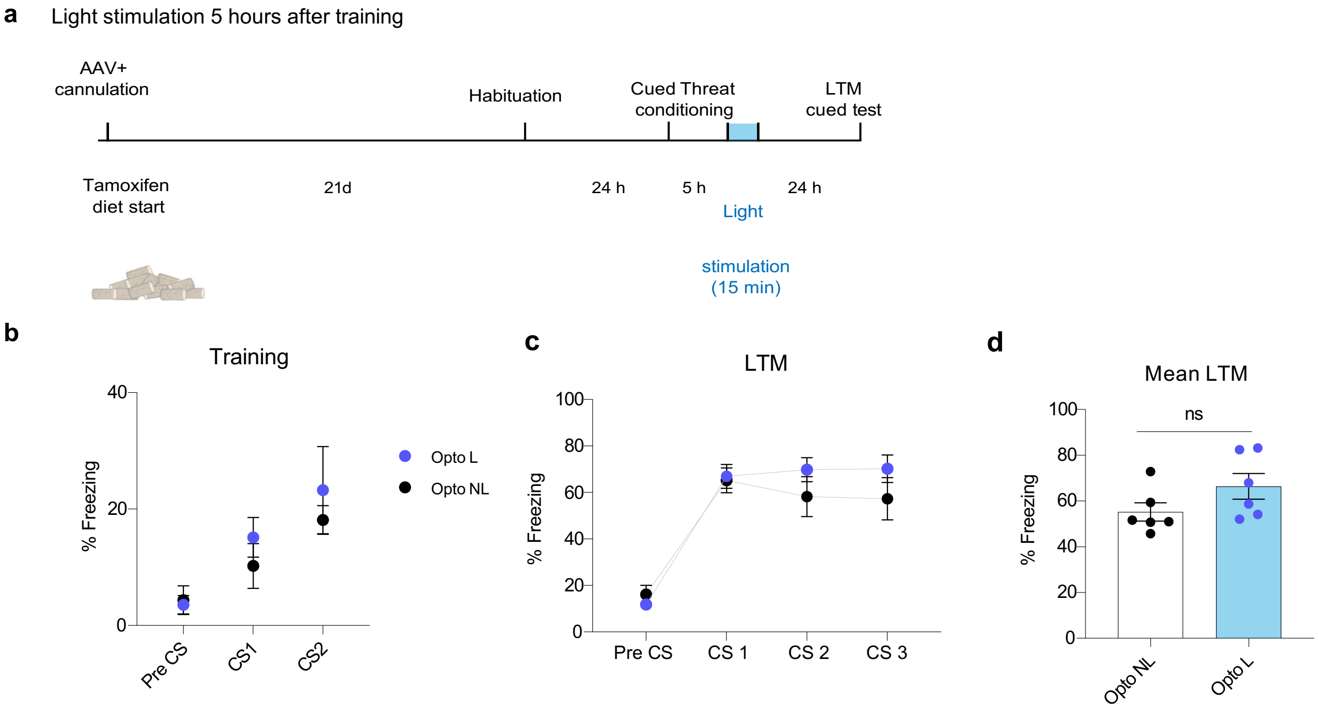


**Supplemental Fig. 4.** Blue light activation of AAV Opto4E-BP 5 hours after cued threat conditioning does not affect LTM formation in mice. a) Light stimulation protocol. CamK2α.Cre $^{ERT2}$ mice infected with AAV Opto4E-BP or mCherry controls in the amygdala underwent cued threat fear conditioning. 5 hours after training they received light stimulation or no light in the amygdala for 15 minutes. 24 hours after training, mice were tested for cued threat LTM. b) Mice learned the association between CS and US during training. RM Two-way ANOVA with Bonferroni's post-hoc test. $F(1,30) = 0.8840$, $p=0.3546$. c) Cued LTM was normal in Opto4E-BP light treated mice compared to controls across all three CS presentations. n=6 per group; RM Two-way ANOVA with Bonferroni's post-hoc test. $F(2,30) = 0.07153$, $p=0.9311$. d) Mean cTC LTM was similar between the two groups. Unpaired Student's t-test ns ($p=0.1424$) n = 6 per group; Unpaired t-test. Data are presented as ± SEM.

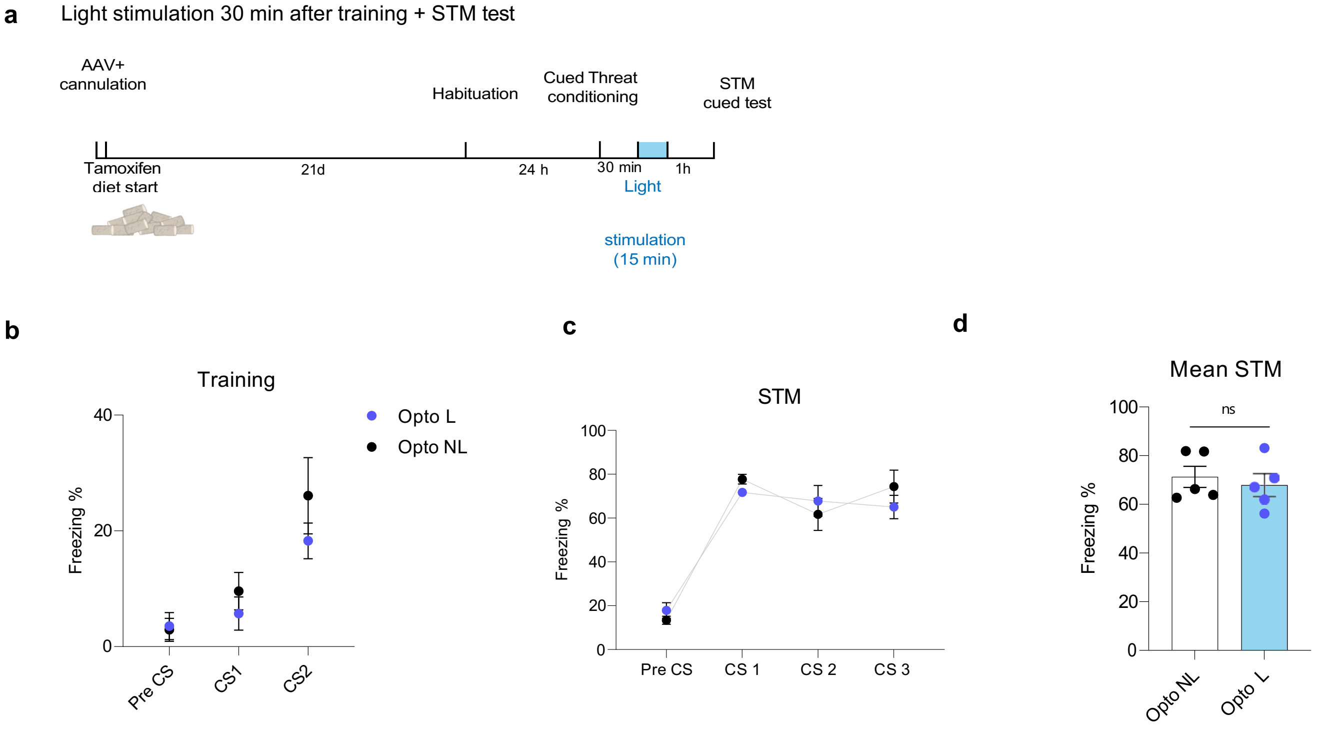


**Supplemental Fig. 5.** Blue light activation of AAV Opto4E-BP 30 minutes after cued threat conditioning does not affect STM formation in mice. a) Light stimulation protocol. CamK2α.Cre $^{ERT2}$ mice infected with AAV Opto4E-BP or mCherry controls in the amygdala underwent cued threat fear conditioning. 30 minutes after training they received light stimulation or no light in the amygdala for 15 minutes. 2 hours after training, mice were tested for cued threat STM. b) Mice learned the association between CS and US during training. RM Two-way ANOVA with Bonferroni's post-hoc test. F (2,27) = 6.1905, p=0.1865. c) Cued STM was normal in Opto4E-BP light treated mice compared to controls across all three CS presentations. n=5 per group; RM Two-way ANOVA with Bonferroni's post-hoc test. F (1,27) = 0.4474, p=0.5092. d) Mean cTC STM was similar between the two groups. Unpaired Student's t-test ns (p=0.5977) n = 5 per group; Unpaired t-test. Data are presented as ± SEM.
